# Laboratory testing of very low-copper-treated water to prolong pupation and emerging time of mosquito larvae: an alternative method to delay mosquito breeding capability

**DOI:** 10.1101/871400

**Authors:** Mohamad Reza, Cimi Ilmiawati

**Affiliations:** Department of Biology, Faculty of Medicine, Andalas Unversity, Padang, West Sumatra, Indonesia; Department of Pharmacology, Faculty of Medicine, Andalas University, Padang, West Sumatra, Indonesia

**Keywords:** *Aedes albopictus*, *Anopheles stephensi*, *Culex pipiens*, copper, larval control, ovitrap, pupation time

## Abstract

**Introduction:** Larvicide application in ovitrap is one of the currently available methods used in mosquito eradication campaign because they eliminate the larval stage. We previously reported that copper in liquid form is a promising candidate due to its potent larvicide properties in a laboratory setting and even in the field. In the field study, several larvae survived in the ovitrap due to the dilution of copper concentration by the rain. The surviving larvae were smaller and less motile. This led our interest to study the effect of a sub-lethal dose of copper in ovitrap on larvae development, pupation time and lifespan in the adult stage.

**Methods:** First instar larvae of three species of mosquito (*Aedes albopictus*, *Culex pipiens* and *Anopheles stephensi*) were bred in copper-treated water at a concentration of 0.60 ppm, 0.30 ppm, and 0.15 ppm and compared with the control group. The surviving larvae were recorded every day in terms of pupa emerging time and adult emerging time. The number of adult mortality was recorded and compared with the control.

**Results:** Copper showed potent larvicide effect in the 0.60 ppm concentration and prolonged pupation time and caused a significantly lower number of emerging mosquitoes down to the lowest concentration of 0.15 ppm. The adult lifespan was not different compared to the control.

**Conclusion:** This study demonstrates the capability of copper below 1 ppm to prolong the pupation time and the emerging time of mosquito larvae. Our findings open the possibility of copper application to cut mosquito breeding capacity that eventually will reduce disease transmission.

## INTRODUCTION

Mosquitoes are causing millions of deaths every year worldwide. More than half of the world’s populations live in areas where mosquitoes are present as vectors and transmit malaria, dengue, Zika, Chikungunya, yellow fever and other diseases [1]. Mosquito-borne diseases are causing an enormous burden on the public health system in many countries with limited financial and human resources. Despite being considered as the most conceivable method to eradicate mosquito-borne diseases [1], global funding for malaria vector control is far below the budget endowed for exploring cures and vaccines for these diseases (US$ 56 *vs* 408 million in 2018) [2]. Lack of funding is considered as one of the challenges in achieving and sustaining malaria control [3].

Core methods for vector control, i.e. insecticide-treated nets (ITNs) and indoor residual spraying (IRS), rely on insecticides. Unfortunately, there is an increase in mosquito resistance to insecticides [4]. Therefore, there is a need to develop novel methods for vector control, particularly practical and economical ones, to be combined with the available methods in integrated vector management. Larval source management (LSM) is part of integrated vector control using strategies to modify or manipulate water bodies as the potential larval habitats of mosquitoes to prevent maturation of mosquito developmental stages. LSM strategies also include the introduction of larvicide (chemical and biological agents) and larvivorous fish (natural predator) into larval habitats [5–7]. Previous studies have proposed the utility of using metallic [8, 9] and liquid [5, 10, 11] copper as a potential and affordable larvicide [7]. Copper is effective as a mosquito larvicide at a concentration below 2 ppm [9], the threshold value deemed safe for human drinking water [12], thus making copper a potential candidate for use in the public health setting.

Our previous findings on the properties of copper to kill mosquito larvae [10] and laboratory testing on the effect of liquid copper at a concentration of 10 ppm on three species of mosquito larvae [11] prompted us to perform a field test in a malaria and dengue-endemic area of West Sumatra, Indonesia. Despite confirming the larvicide effect of copper [7], we observed that dilution by rainwater should be considered as confounding in the field since we noted that some larvae survived in outdoor ovitraps. However, the survived larvae appeared to be less motile and were smaller in size compared to unexposed larvae. Consequently, further study is required to determine whether the survived larvae capable to continue their life cycle until adulthood. Therefore, we performed a laboratory test of sub-lethal concentrations of copper exposure to mosquito larvae to verify its effect on mosquito development and breeding capacity.

## MATERIALS AND METHODS

### Ethical Approval

This research was approved by the ethics committee of the Faculty of Medicine, Universitas Andalas (Approval No.036/KEP/FK/2019).

### Mosquito Eggs and Mice

*Anopheles stephensi* (strain SDA 500), were reared in our laboratory under room temperature (26°C), 50-70% relative humidity, and a 13:11 h light: dark cycle. From the day of emergence, mosquitoes were provided with a 5% fructose solution soaked in a filter paper. Females five days after emergence were allowed to feed on anesthetized mice. Three days later, an oviposition dish was placed in a cage containing gravid females. The eggs were laid on a filter paper soaking in the dish. The filter paper with eggs was placed on a 12 × 20 cm hatching tray containing 500 ml of water. After hatching, 3-5 mg of carp food per tray was sprinkled on the surface of the water twice daily. Twelve days later, pupae were collected daily and transferred to cages for adult emergence.

Eggs of *Aedes albopictus* and *Culex pipiens* were purchased and the first instar newly hatched larvae were utilized in the experiment. Female BALB/c mice were purchased from SLC (Shizuoka, Japan). The mice were fed *ad libitum* and exposed to a 13:11 h of light: dark cycle. This mosquito-mouse cycle was used to maintain mosquitoes in the laboratory.

### Preparation of Copper Sulfate Solutions

Four concentrations of CuSO_4_ solution were used for the experiment. The copper concentrations were 0.60, 0.30, 0.15, and 0 ppm. Each solution was allocated to a pre-washed square plastic plate and covered with a transparent plastic plate. First, we prepared a standard solution of 100 mM of CuSO_4_ by mixing copper sulfate powder with filtered water (Fine Ceramic Filter NGK insulators, Nagoya, Japan). Then, we prepared the four concentrations for the experiment by diluting them from the standard solution and confirming the copper level using a copper measuring device (Hanna Instruments, Tokyo, Japan) and Z-5010 Polarized Zeeman flame atomic absorption spectrophotometer (Hitachi Ltd, Tokyo, Japan).

### Preparation of Mosquito Larvae

First instar newly hatched larvae were used in this experiment. Fifty larvae were separated and put inside each container. Three containers for each concentration of copper were prepared. Larvae were allowed to hatch and were observed every 24-hours under the microscope for mortality. A fine ceramic filter (NGK Insulators, Nagoya, Japan) was used to remove chlorine ions from tap water for the experiment.

## RESULTS

To determine the effect of very low concentrations of copper on mosquito larval mortality, larvae of *A. albopictus*, *A. stephensi* and *C. pipiens* were exposed to 0.60, 0.30 and 0.15 ppm of copper solutions (CuSO_4_) and were observed daily. Larvae of *A. stephensi* and *C. pipiens* exposed to 0.60 ppm of copper showed 100% mortality within a week, while a small number of *A. albopictus* larvae survived. Larvae from three species roughly showed 50% mortality when exposed to 0.30 ppm of copper for seven days. CuSO_4_ at 0.15 ppm only had a statistically significant effect on *C. pipiens* larval mortality compared to control, started on the third day of exposure (**Figure 1**).

**Figure 1.**
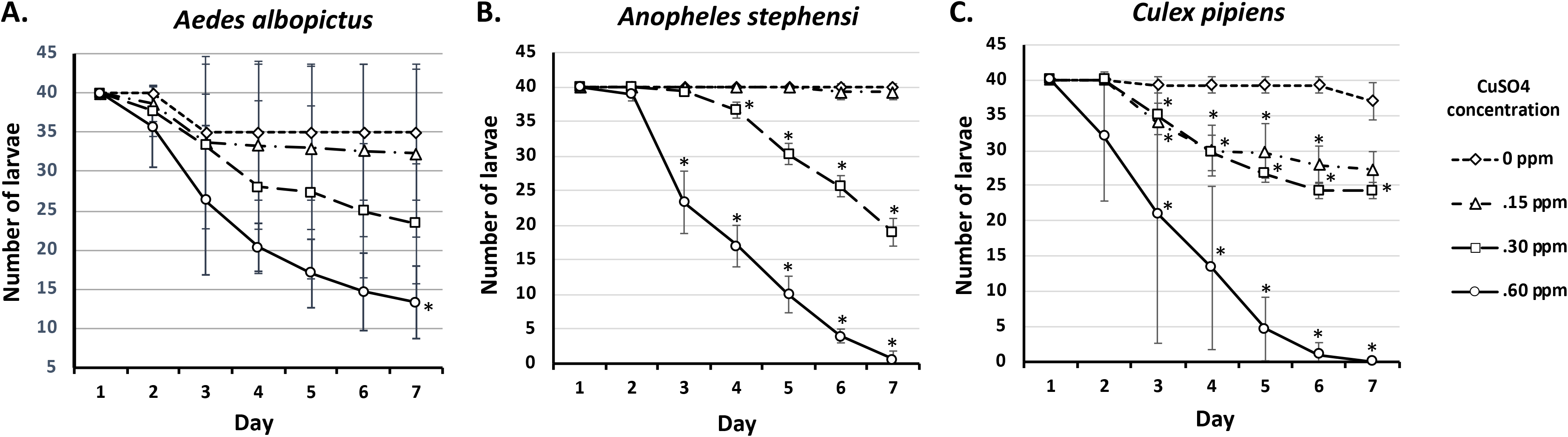
The observed mortality of *A. albopictus*, *A. stephensi*, and *C. pipiens* larvae on exposure to 0.15, 0.30, and 0.60 ppm of CuSO_4_. *significantly different from control, p<0.05 (t-test or Mann-Whitney U test)

To figure out the effect of very low concentrations of copper exposure to larval pupation time, the number of emerging pupae from the three species were counted. Pupae started to emerge at day eight to 10, depending on the mosquito species. Surviving larvae exposed to 0.15 and 0.30 ppm of copper showed prolonged pupation time in all species observed compared to the control group. We also observed that the surviving larvae of *Ae. albopictus* on 0.60 ppm were not turned into pupae (**Figure 2**).

**Figure 2.**
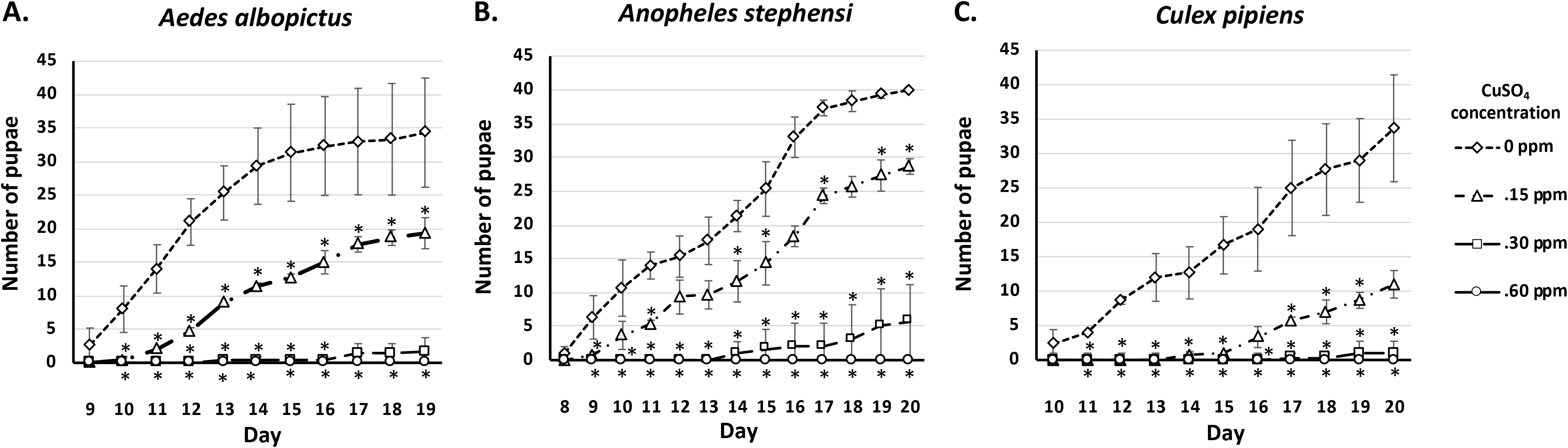
The number of emerging pupae of *A. albopictus*, *A. stephensi*, and *C. pipiens* on exposure to 0.15, 0.30, and 0.60 ppm of CuSO_4_. *significantly different from control, p<0.05 (Mann-Whitney U test)

To determine the effect of very low concentrations of copper exposure on mosquito ability to reach adulthood, we observed and counted the number of emerging adults in three exposed species. Surviving pupae of all observed species exposed to 0.15 ppm of CuSO_4_ become adult mosquitoes, albeit at a statistically significantly lower number compared to the control group. Adult mosquito of all observed species emerged at later days compared to control and survived up to 22 days of observation. Almost none of the pupae of all observed species exposed to ppm of CuSO_4_ emerged as an adult mosquito (**Figure 3**).

**Figure 3.**
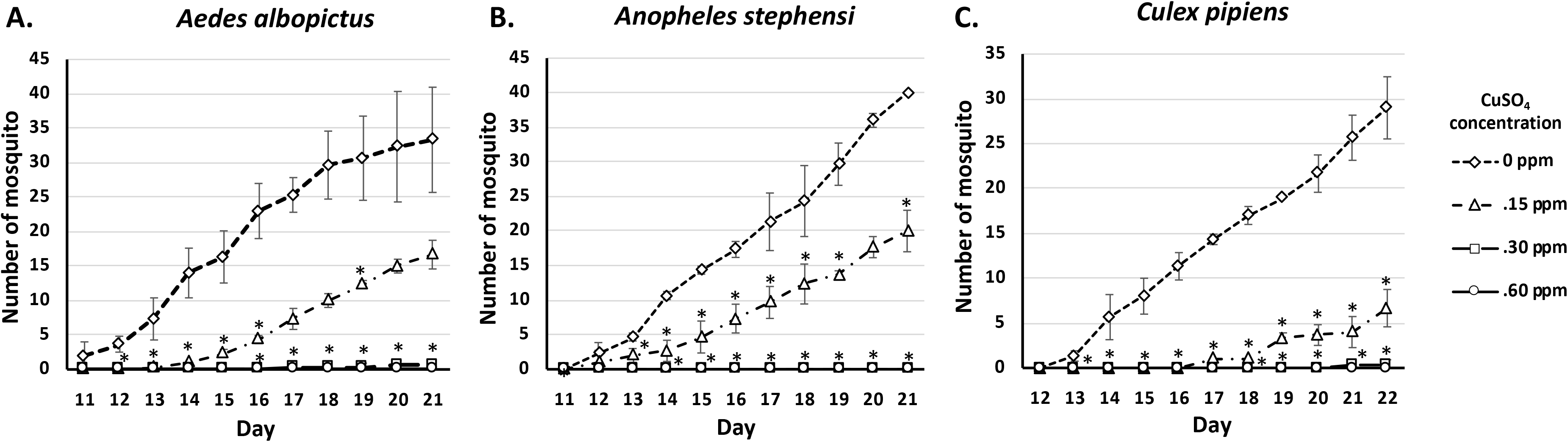
The number of emerging *A. albopictus*, *A. stephensi*, and *C. pipiens* mosquito on exposure to 0.15, 0.30, and 0.60 ppm of CuSO_4_. *significantly different from control, p<0.05 (Mann-Whitney U test)

It is noteworthy that all adults emerged in copper-treated groups and control appeared to be viable and showed no difference in term of their lifespan. It seemed that the copper effect on the survived larvae was limited to prolonging the emergence of pupae and was not carried on to the adult stage.

## DISCUSSION

The results of the present study demonstrate that copper adequately lead to a high rate of larval mortality in at a concentration below 1 ppm (0.60 ppm). At a further lower concentration (0.30 ppm), copper still kill half of the larvae, and together with the lowest concentration of 0.15 ppm, prolongs pupation time and the emerging time of adult mosquitoes compared to the control group. The number of mosquitoes that emerged afterward is also significantly lower than those in the control group.

However, the copper effect is limited in the larvae and pupae stage. Copper shows no effect on the adult stage. The effect on the larvae probably caused by the copper effect to commensal bacteria in the larval midgut, which disrupts their intestinal function [13]. This prevents them to eat properly and to collect enough energy for pupation and emerge as an adult mosquito. Once the larvae become adult, this copper effect no longer plays a role in their life cycle.

This is an interesting finding, since at a very low concentration copper prolongs breeding time of mosquito. This might lead to the possibility to utilize copper in a tap water to jeopardize mosquitoes breeding capability. The mosquitoes will need a longer time to breed, which then reduce their capability to transmit mosquito-borne diseases such as dengue hemorrhagic fever.

It is also known that *Aedes sp.* are resistant to insecticide in indoor or outdoor spraying. A study conducted in Sri Lanka showed *Aedes aegypti* and *Aedes albopictus* were highly resistant to DDT and they can oviposit indoors and outdoors [14]. Another study in Bangladesh reported abundant potential larval habitats for *Aedes sp.* in containers or jars spread around the city [15]. These situations need another approach of vector control, such as laying ovitraps or another effort to cut or delay mosquito breeding capability, or even provide insecticide in the tap water if possible.

The US EPA is suggesting 1.3 ppm as a safe limit of copper concentration in the drinking water [16], thus making the application of very low 0.15 ppm of copper might be logically acceptable in the future.

We believed that, together with the application of copper in the ovitraps, the concentration of 0.15 ppm of copper in the tap water in houses and large scale water reservoir might be a potential strategy in cutting mosquito breeding capability and their ability to transmit mosquito-borne diseases, especially dengue hemorrhagic fever, which mainly breed in the city water reservoir and container. This result might be a chance to re-evaluate the allowance of copper in drinking water as one of the possible strategies in eradicating mosquito-borne diseases in the future.

## Authors’ contributions

MR conceived and designed the study protocol. MR and CI formulated the study proposal and obtained funding. MR conducted research in the laboratory. MR carried out data analysis and interpretation and wrote the manuscript, while CI conducted the statistical analysis. All authors read and approved the final manuscript.

## Acknowledgment

The authors thank Prof. Hirotomo Kato, Ph.D for his kind permit to use his laboratory in Japan for this research.

## Conflict of interest

The authors report no conflicts of interest.

## Notes

#### Summary of Updates

The Introduction part has been revised to avoid similarity with the author's previous published papers.

